# Diet induces parallel changes to the gut microbiota and problem solving performance in a wild bird

**DOI:** 10.1101/827741

**Authors:** Gabrielle L. Davidson, Niamh Wiley, Amy C. Cooke, Crystal N. Johnson, Fiona Fouhy, Michael S. Reichert, Iván de la Hera, Jodie M.S. Crane, Ipek G. Kulahci, R. Paul Ross, Catherine Stanton, John L. Quinn

## Abstract

The microbial community in the gut is influenced by environmental factors, especially diet, which can moderate host behaviour through the microbiome-gut-brain axis. However, the ecological relevance of microbiome-mediated behavioural plasticity in wild animals is unknown. We presented wild-caught great tits (*Parus major*) with a problem-solving task and showed that performance was weakly associated with variation in the gut microbiome. We then manipulated the gut microbiome by feeding birds one of two diets that differed in their relative levels of fat, protein and fibre content: an insect diet (low content), or a seed diet (high content). Microbial communities were less diverse among individuals given the insect compared to those on the seed diet. Individuals were less likely to problem-solve after being given the insect diet, and the same microbiota metrics that were altered as a consequence of diet were also those that correlated with variation in problem solving performance. Although the effect on problem-solving behaviour could have been caused by motivational or nutritional differences between our treatments, our results nevertheless raise the possibility that dietary induced changes in the gut microbiota could be an important mechanism underlying individual behavioural plasticity in wild populations.

## INTRODUCTION

The enteric microbial community, frequently referred to as the gut microbiome, is an important ecosystem that contributes to host behaviour ^1,2^. Recent evidence points to a communication link between the host’s gut microbiome and the brain, known as the microbiome-gut-brain axis ^3,4^. Experimental alteration of the gut microbiome can impact important behaviours and cognition, including learning, memory, anxiety, activity levels and social interactions ^3,5–8^, as well as cause changes to neurogenesis ^9^ and protein expression in the brain ^7,10,11^. Neurotransmitters and short chain fatty acids released by microbes act as signals that can be communicated to the brain ^1^ and the extent to which signalling occurs is dependent on the microbial taxa present ^12^. Evidence of the microbiome-gut-brain axis affecting behaviour is limited to experiments on model laboratory animals and to correlational studies on mental health in humans ^1,4^. Little is known about whether these findings can be applied to natural populations, where microbiome-host interactions are predicted to have important effects on traits that directly impact animal fitness ^13,14^, such as cognition ^15^ and foraging behaviour ^16^.

Several environmental factors have been shown to contribute to enteric microbial community composition in wild vertebrates, including geographical location ^17,18^, habitat characteristics ^19,20^, and seasonality ^21–23^. These temporal and spatial differences in the microbiome, between and within populations, also correlate with variation in diet ^21,24^. Reports of direct effects of dietary manipulations on the gut microbiota in birds to date have been largely restricted to poultry ^25^, and a recent study in wild-caught house sparrows demonstrated diversity and family-level changes to microbiota as a consequence of dietary manipulation ^26^. Changes in diet can also lead to a significant alteration to the gut microbiome within 24 hours in humans ^27^, where changes are dependent on dietary features such as protein levels ^27,28^, fat levels ^29^, and whether plants are present in the diet ^30,31^. Youngblut et al. ^32^ found diet explained a high proportion of variation in gut microbiota across vertebrate species including a subset of avian species. Hird et al ^33^ also showed that variation in gut microbiome across species was explained by diet in neotropical birds. Thus, microbiome plasticity should be particularly important in wild populations ^13,14^, many of which are subject to regular changes in food availability ^34^, and in which individuals typically differ in their foraging strategies ^35^. Studies in model laboratory organisms show that phenotypic plasticity in cognitive performance and anxiety-like behaviour can occur in parallel with gut microbiome alterations caused by diet ^8,36,37^. For example mice fed beef-chow had a higher microbial diversity than those fed on normal chow (diets which differed in nutritional, amino acid and fatty acid content) and showed improved working and reference memory ^36^. Observational studies also point to a relationship between diet, cognition and mood in humans ^38,39^, effects that may be mediated by the host gut microbiome ^39^. Although some empirical studies have been published on lab model systems, there is now a need for manipulative experiments using model ecological organisms to understand how dietary induced changes in the microbiome might affect functionally significant behaviours in the wild ^13^.

The great tit (*Parus major*) is a highly innovative species that has long been a model organism in field studies on behaviour, ecology and evolution. It is also ideal for short-term laboratory studies because they adapt well to temporary captivity ^40,41; G. Davidson personal observation^. Innovative problem solving performance’ (PSP), in which individuals must solve a technical problem that they have not encountered previously to obtain a reward, is often described as being underpinned by cognitive mechanisms, as well as other factors such as motivation, persistence, exploration ^42^ and parasite infection ^40^. In wild great tits, PSP differed consistently over time and across different tasks among individuals ^43^. However, having solved this task previously in a captive environment did not predict whether the same bird solved the same task in the wild ^44^, suggesting that despite consistent individual differences, PSP is also highly plastic. Quantitative genetic analysis in one population suggested that adult PSP had little if any heritable component and instead was primarily linked to natal habitat characteristics associated with suitable food items for nestlings ^45^. PSP in the great tit has been linked to a range of behaviours and fitness-related traits in the great tit, both in the wild and in the laboratory ^40,41,45^. More generally, innovativeness is positively correlated with dietary breadth across bird species ^46^ because innovations increase their access to a wide range of resources ^47^.

Here we tested whether diet and habitat predicted gut microbiota metrics in wild-caught great tits, and whether natural and manipulated gut microbiome profiles correlated with PSP. First we sampled birds from both rural and urban areas, known to differ in their quality as habitat for great tits ^e.g. 48^, and therefore we expected host gut microbiota may differ in diversity, community structure (i.e. beta diversity) and/or taxa-level abundance ^e.g. 49^, though our replication at the site level was limited for each habitat. Alpha diversity measures the number of different species and depending on the type of metric, can also take into account how evenly different taxa are dispersed within the microbial community. Beta diversity looks at differences in the presence and absence of taxa between groups. Taxa-level abundance measures the proportion of a taxonomic group relative to the rest of the community. In line with our hypotheses above, we predicted a positive correlation between PSP and natural microbiome alpha diversity, although we also tested for differences in microbial community structure similarity/dissimilarity (i.e. beta diversity) and taxa-level relative abundance between solvers and non-solvers. Second, we manipulated the diet of wild-caught great tits to test whether microbiota community structures became more dissimilar, and whether phylum-level and genus-level taxa changed within and between treatment groups. Given the seed diet was higher in fat, protein and fibre content than the insect only diet, we expected these individuals to have higher alpha diversity, higher Firmicutes and lower Bacteroidetes if similar dietary effects occur in great tits as have been reported in humans ^e.g. 28^. Third, we tested whether a change in the gut microbiome as a consequence of the manipulated diet influenced PSP by presenting these birds with the same task again. We predicted that if the gut microbiome influenced host PSP, rather than the dietary treatment itself, then a change in PSP should specifically be associated with the same metrics of the gut microbiome as those that were changed as a result of dietary manipulation, whilst controlling for the dietary treatment.

## RESULTS

### Sequencing summary

We collected fresh faecal samples within one hour of arrival in captivity, and 13 days later at the end of the experimental treatments, to quantify the gut microbial community profiles using 16S RNA sequencing. We obtained 4,109,746 sequences distributed along 2306 OTUs with an average 67,373 ± 4,873 se sequences per sample (min. 16,126 and max. 180,097). A total of 314 OTUs from 192 genera, 116 family and 19 phyla were present in the data, following filtering steps.

### Relative abundance, diet and microbial diversity

The most prominent phyla across all samples were as follows (mean percentage relative abundance ±SE): Proteobacteria (55.3%±4.0); Cyanobacteria (14.8%±2.6); Firmicutes (10.2%±2.1); Tenericutes (9.0%±2.3); Actinobacteria (4.0%±1) and Bacteroidetes (2.0%±0.5). Mean relative abundance of most abundance phyla across treatment groups is depicted in Figure 1 and Table 1, and all phyla per sample (Figure 2.). Proteobacteria increased significantly over the course of captivity following the insect diet (z=2.02, p=0.04), but not in the seed diet (z=-0.28, p=0.78), and tended to be higher in adults than juveniles (z=1.84, p=0.07). Bacteroidetes (natural log (Ln)-transformed) increased following the insect diet (t=22.22, p=0.03), but not the seed diet (t=1.01, p=0.32), and tended to be higher in urban compared to rural habitats (t=1.80, p=0.08). Actinobacteria was significantly higher in birds from urban habitats (t= 2.47, p=0.02). There was no significant effect of diet, habitat, sex or age on Firmicutes (Ln-transformed) or Tenericutes (Ln-transformed) (Table 2). There were no significant interactions between diet and habitat for all models. Full model outputs are provided in Table S2, supplementary.

**Table 1.** Relative abundance of the most abundant phyla at Day 1 (pooled data of all Day 1 samples), Day 1 for birds assigned the insect diet (ID), Day 1 for birds assigned the seed diet (SD), and birds after the insect diet and the seed diet. Figures represent means and standard errors.

| Phylum | Day 1 | Day 1 (ID) | Day1 (SD) | Insect diet | Seed diet |
| --- | --- | --- | --- | --- | --- |
| Proteobacteria | 51.0%±5.3 | 50.9%±7.3 | 51.0%±8.0 | 76.0±7.9 | 53.8%±8.3 |
| Cyanobacteria | 21.0%±4.0 | 14.1%±3.1 | 29.6%±7.8 | 0.3%±0.1 | 14.5%±4.6 |
| Firmicutes | 9.4%±2.0 | 10.5%±3.5 | 8.2%±0.03 | 11%±7.1 | 10.4%±4.5 |
| Tenericutes | 7.8%±3.3 | 10.7%±5.7 | 4.2%±2.7 | 7.0±3.1 | 15.8%±5.8 |
| Actinobacteria | 3.5%±1.3 | 3.4%±2.1 | 3.6%±1.5 | 2.1%±1.0 | 1.3%±0.3 |
| Bacteroidetes | 0.5%±0.1 | 0.5%±0.1 | 0.4%±0.2 | 2.3%±0.8 | 1.6%±1.1 |
| Others | 6.6%±0.2 | 9.8%±6.1 | 2.9%±2.1 | 1.4%±0.1 | 2.6%±1.1 |

**Table 2.** Summary of the relationships between microbiota and host traits. Significant relationships are shown as either positive (+), negative (-) or as p-values (i.e. beta-diversity where differences do not represent directionality). NS = Non-significant.

| Microbiota metrics | Diet | PSP | Habitat | Sex | Age |
| --- | --- | --- | --- | --- | --- |
| **Proteobacteria** | (+) |  |  |  |  |
|  | insect | NS | NS | NS | NS |
| **Firmicutes** | NS | NS | NS | NS | NS |
| **Bacteroidetes** | (+) |  |  |  |  |
|  | insect | NS | NS | NS | NS |
| **Actinobacteria** |  |  | (+) |  |  |
|  | NS | NS | urban | NS | NS |
| **Tenericutes** | NS | NS | NS | NS | NS |
| **Chao1** | (-) |  |  |  |  |
|  | insect | (+) | NS | NS | NS |
| **Shannon's index** | NS | (+) | NS | NS | NS |
| **Observed OTUs** | NS | (-) | NS | NS | NS |
| **Jaccard** | p < 0.01 | NS | p < 0.01 | NS | NS |
| **Weighted unifracs** | p < 0.01 | NS | p = 0.01 | NS | p < 0.05 |
| **unweighted unifracs** | p < 0.01 | p < 0.05 | NS | NS | NS |

**Figure 1.**
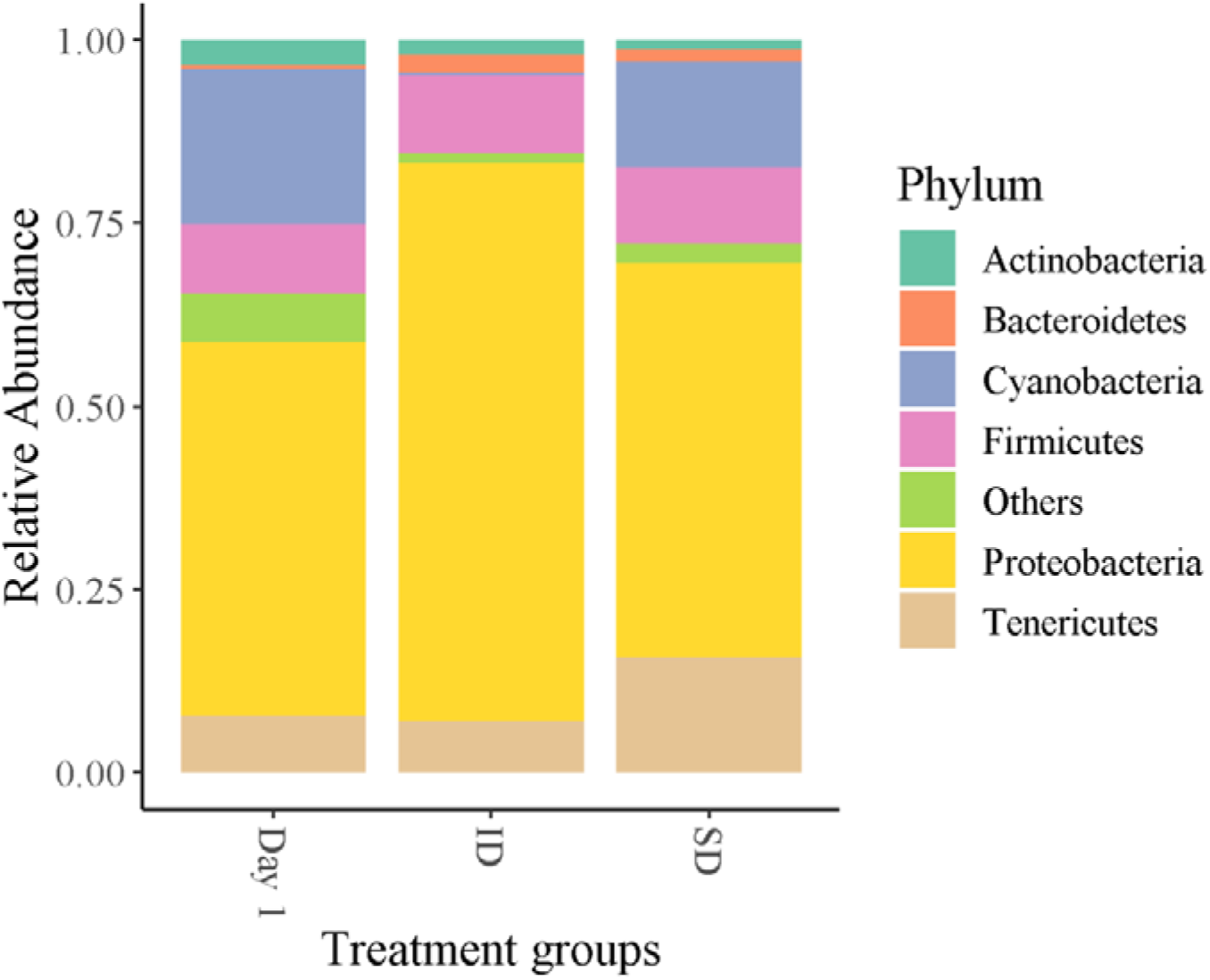
Relative abundance of top seven phyla across dietary treatment groups. Proteobacteria and Bacteroidetes significantly increased following the insect diet, but not the seed diet.

**Figure 2.**
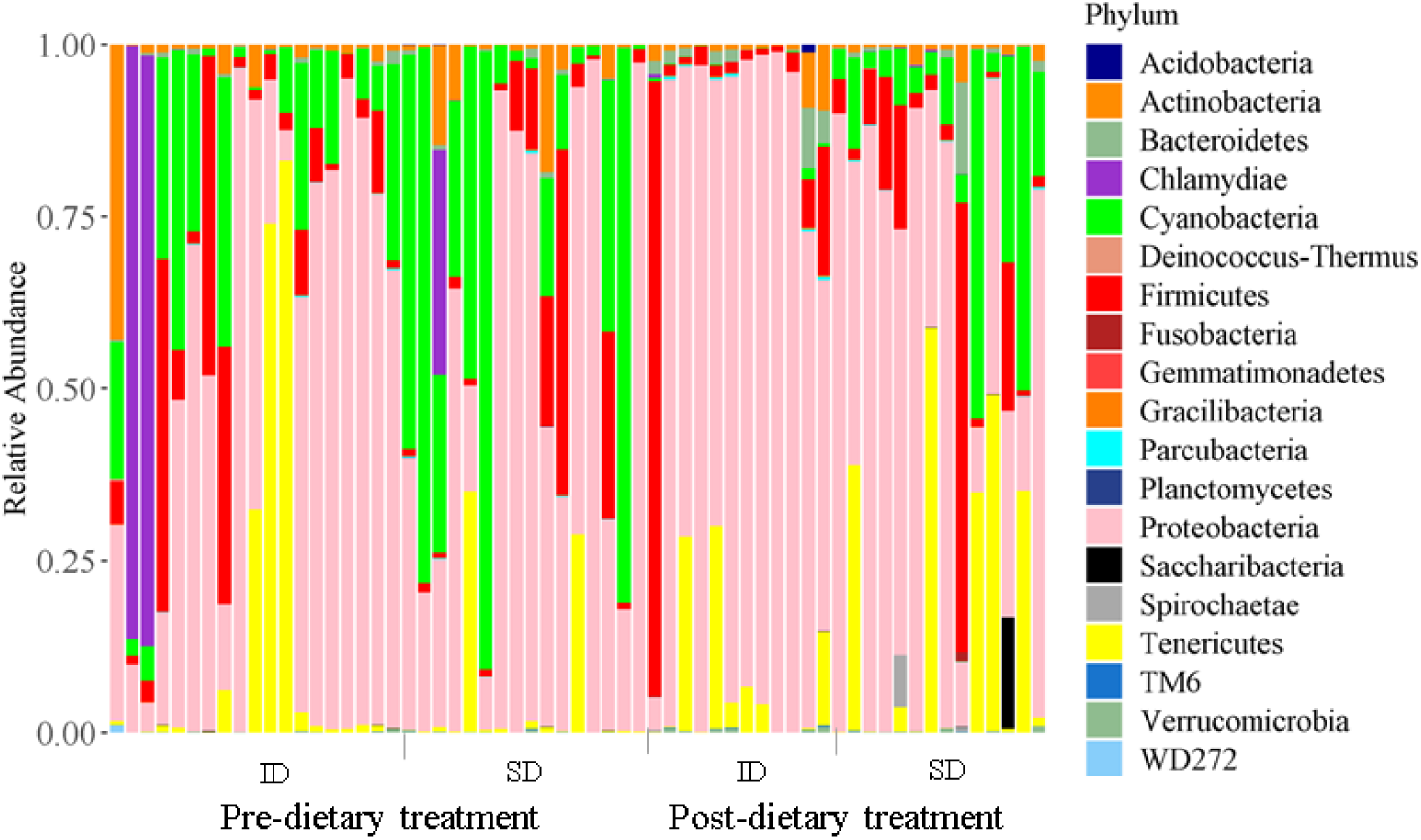
Relative abundance of bacterial phyla per sample for birds before and after their dietary assignments of either insect diet (ID) or seed diet (SD).

Significant differences in genus-level abundance attributed to dietary treatments were found for 22 genera. Birds given the insect diet showed a decrease in *Devosia, Rickettsiella, Sphingomonas, Pantoea, Arthrobacter, Brevibacterium, Brachybacterium, Clostridium* and *Carnobacterium*, and an increase in *Candidatus and Methylobacterium*. Birds given the seed diet showed a decrease in *Cronobacter* and *Serratia*, and an increase in *Microbacterium*. Birds in both dietary groups showed a decrease in *Bradyrhizobium, Staphylococcus* and *Rahnella*, and an increase in *Lactobacillus, Bacillus, Ureaplasma, Delftia, Flavobacterium, Streptococcus* and *Rhodococcus* (Table S3, supplementary).

There was a significant decrease in alpha diversity, specifically Chao1 (square root transformed) following the insect diet (t=-2.51, p=0.02), but not the seed diet (t=-0.07, p=0.94) (Figure 3a). A similar pattern was found in Shannon’s index, although the effect of the insect diet was marginally non-significant (insect diet: t=-2.02, p=0.06; seed diet: t=0.01, p=0.99) (Table 2, Figure 3b). Number of observed OTUs (Ln-transformed) did not differ significantly across treatments (insect diet t=-1.74, p=0.10; seed diet t=0.43, p=0.67) (Figure 3c). There was no significant effect of sex, age, or habitat type on alpha diversity. There was no significant interaction between diet and habitat. (Table S4, supplementary). ADONIS tests showed dietary treatment significantly influenced beta diversity across all three metrics (Table 2, Figure 4a and Table S5). Post-hoc tests with Bonferroni correction indicate that the differences were between day 1 and the post-diet insect group (R2=0.07, p<0.01), and between the two post-diet groups (R2=0.10, p=0.03). Beta diversity also significantly differed between habitat types (Table 2, Table S5).

**Figure 3.**
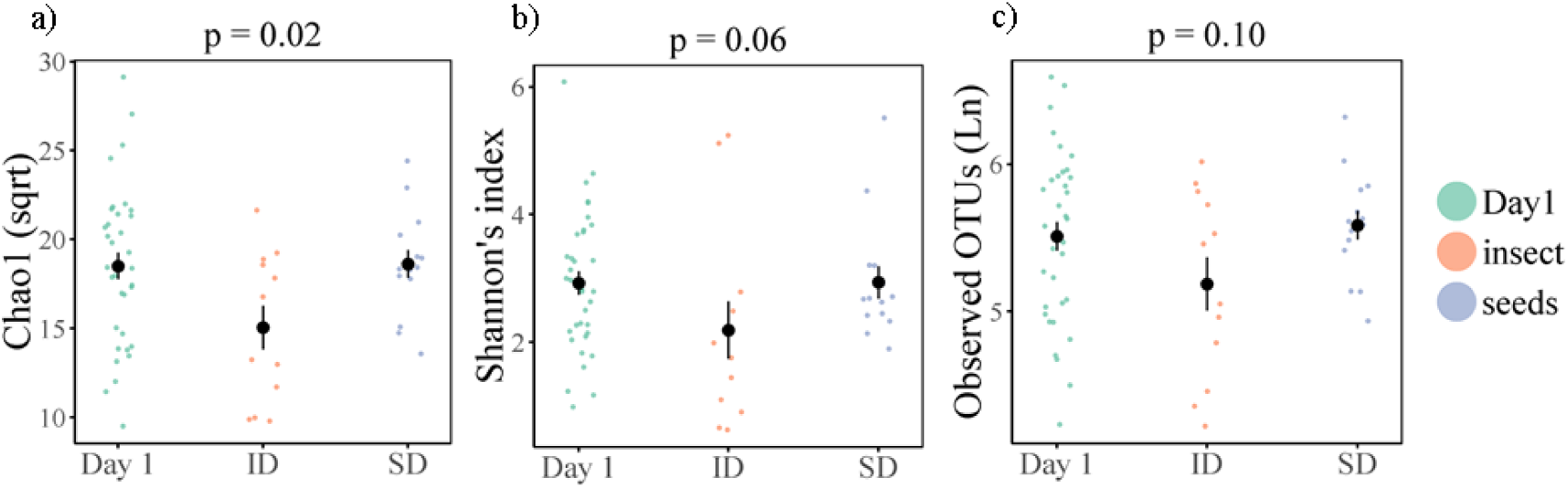
Alpha diversity for birds pre-dietary assignment (Day 1), and following the insect diet (ID) and the seed diet (SD) for a) Chao1 (sqrt), b) Shannon’s index, c) Observed OTUs (Ln). Coloured points denote individual data points, black points and line denote mean and ± SE. p-values represent the comparison between post-insect diet group and day 1.

**Figure 4.**
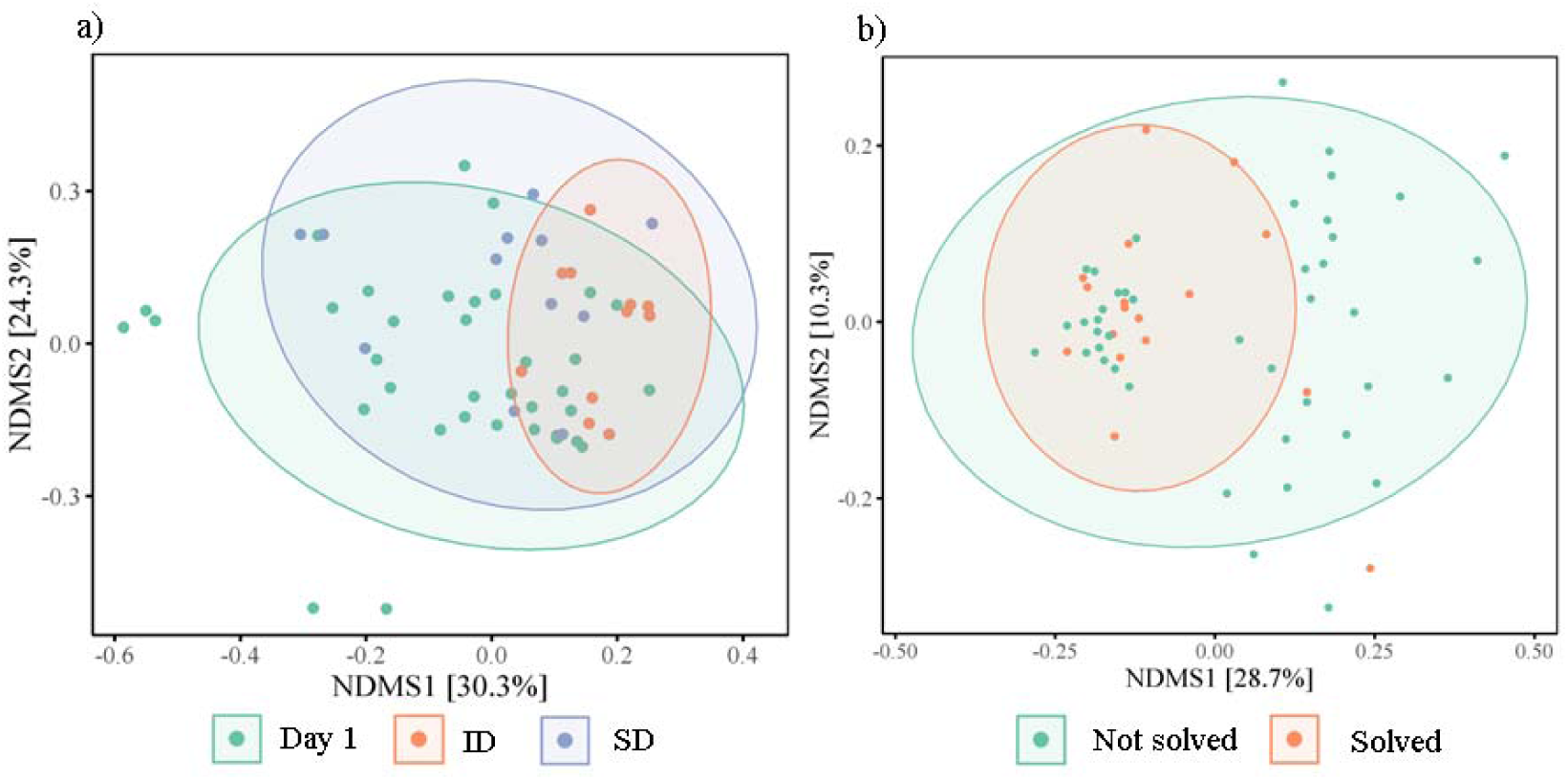
Nonmetric Multidimensional Scaling (NMDS) ordination plots derived from beta diversity measurements for (a) weighted unifrac distances of diet (Day 1, insect diet (ID) and seed diet (SD), and (b) unweighted unifrac distances of problem solving performance (not solved, solved). Ellipses represent standard deviations around the centroids of the groups. Numbers in brackets refer to the variance explained by NDMS axes.

### Problem solving, diet and microbial diversity

To quantify individual innovative problem solving performance, naïve birds were presented with a foraging task, in which birds could either pull a stick to release a platform holding a food reward (wax moth larvae), push a door to gain access to the reward, or pull a string attached to the reward. The PSP task was solved 17 times, by 15 different individuals across both trial days, with two individuals solving both before and after the dietary treatment. Twelve birds solved on day 1 (4 lever, 8 door), 5 birds in the seed group solved (2 lever, 3 door), and none of the birds from the insect group (post-diet treatment) solved. Beta diversity prior to dietary manipulation tended to differ between problem solvers and non-problem solvers (unweighted unifrac distances R = 0.05, p=0.07) (Figure 5a), and birds with lower alpha diversity were less likely to solve, although this was non-significant (z=-1.54, p=0.13) (Figure 5b). All other metrics of natural variation in the gut microbiome were not significantly associated with PSP (Table S4). However, PSP across both trials, when controlling for dietary treatment and repeated measures, was positively correlated with alpha diversity (Shannon: z=2.22, p=0.03; Chao1 z=2.13, p=0.04, observed OTUs z=1.96; p=0.06) (Table S2, Figure 6a,b,c), and beta diversity (unweighted unifrac: R2=0.02, p=0.04) (Table 2, Table S5, Figure 4b). Phylum-level and genus-level relative abundance was not associated with PSP.

**Figure 5.**
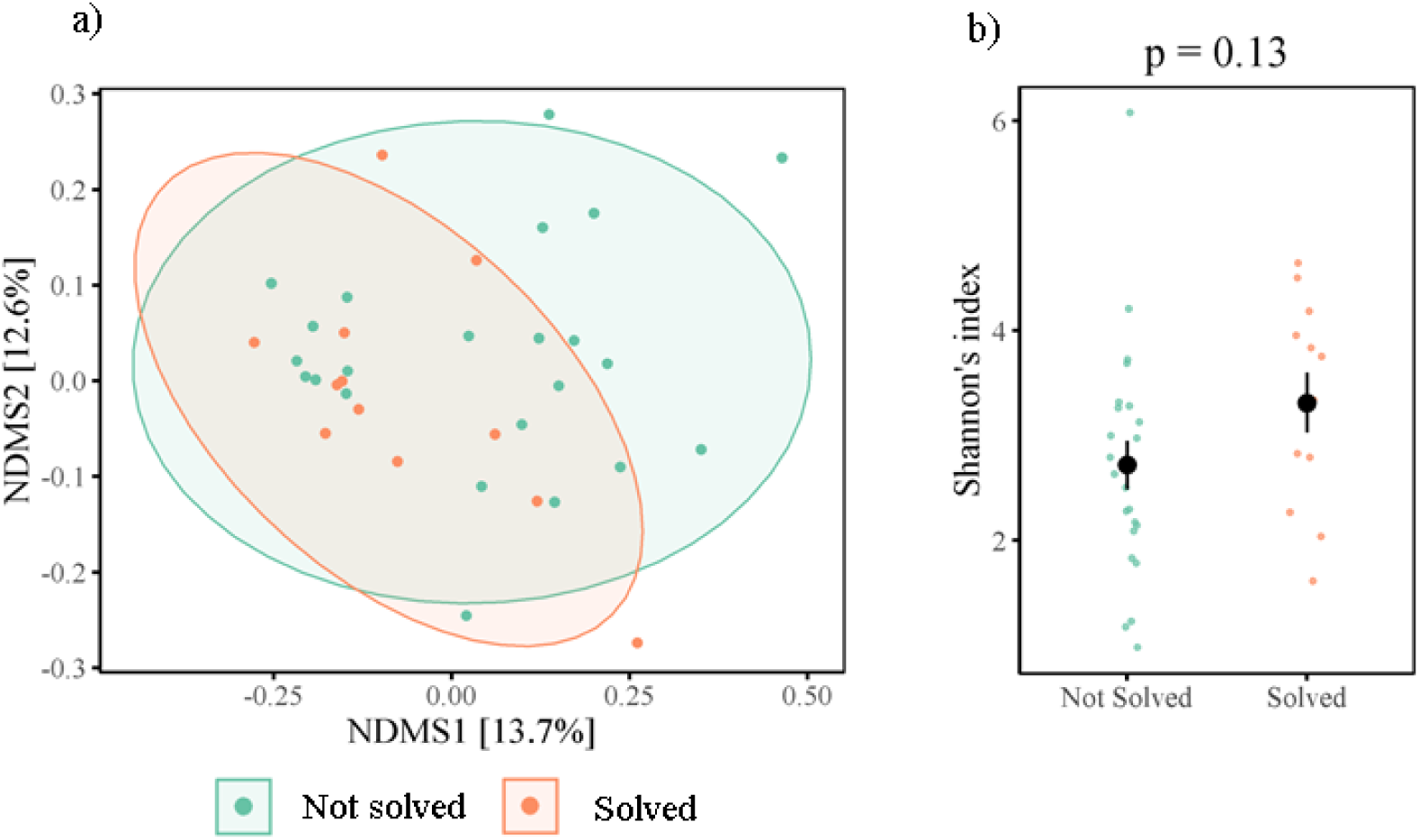
Natural variation in gut microbiome diversity and problem solving performance for a) beta diversity (Nonmetric Multidimensional Scaling (NMDS) ordination plots), and b) Shannon’s index. Coloured points denote individual data points, black points and line denote mean and ± SE.

**Figure 6.**
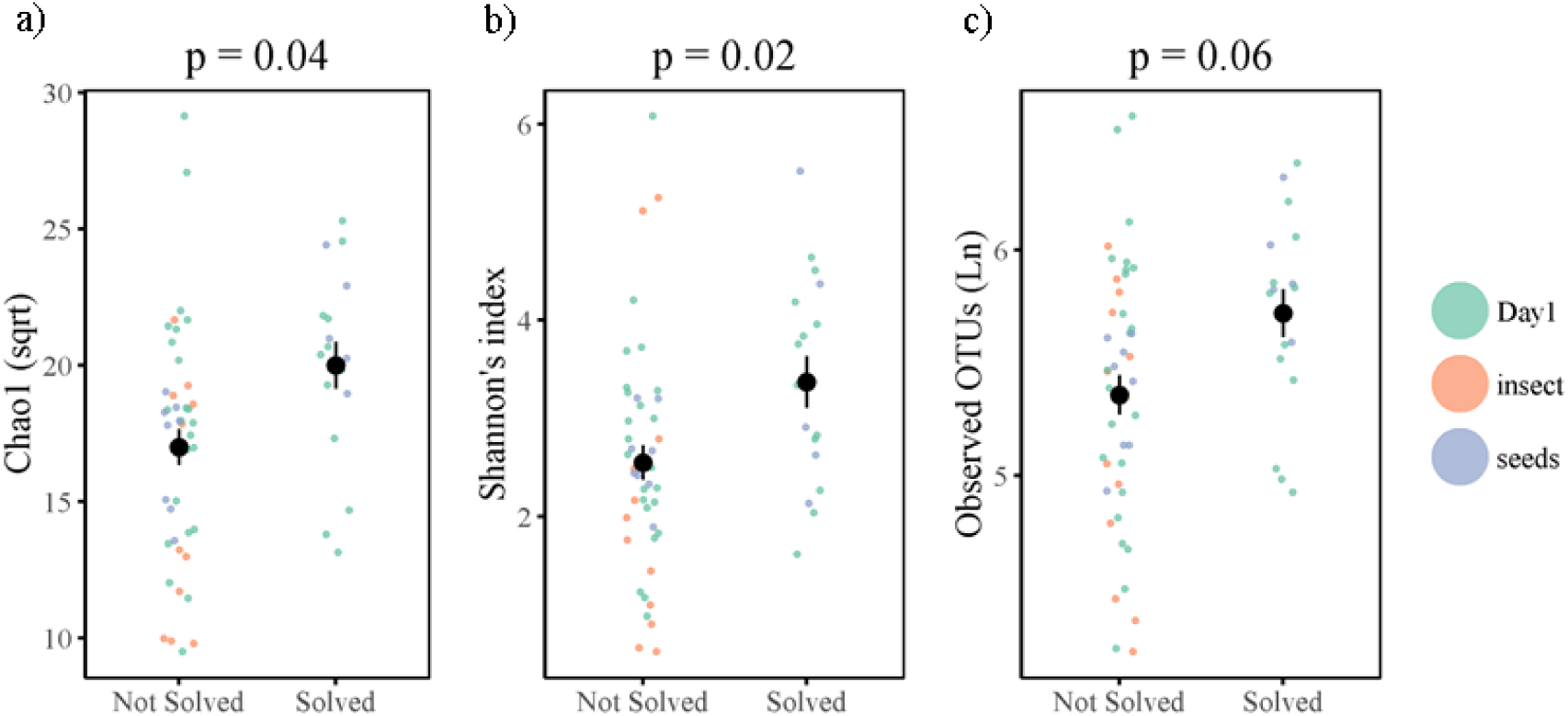
PSP-alpha diversity relationships in a) Chao1 (sqrt), b) Shannon’s index, c) Observed OTUs (Ln) including data points from both day 1 and day 12. Coloured points denote individual data points, black points and line denote mean and ± SE.

Birds assigned to the seed diet solved more than the birds assigned to the insect diet (z=2.22, p=0.03), and birds tended to be more likely to solve on day 1 than day 12 (z=1.93, p=0.054), though this effect was likely driven by birds assigned to the insect diet (Figure 7). There was a tendency for juveniles to solve more than adults (z=1.94, p=0.053). Neither the interactions between habitat, nor between experiment day and diet were significant (Table S2, supplementary). PSP was not attributed to differences in motivation to consume the food reward as all birds were equally likely to consume the same reward when it was made freely available to them, regardless of dietary treatment and experiment day (diet z=1.18, p=0.24, experiment day z=0.85, p=0.40). To rule out effects of glucocorticoids on cognitive performance [47], we found that performance was not attributed to individual baseline or stress-induced blood-circulating corticosterone levels measured as part of a separate experiment (unpublished data).

**Figure 7.**
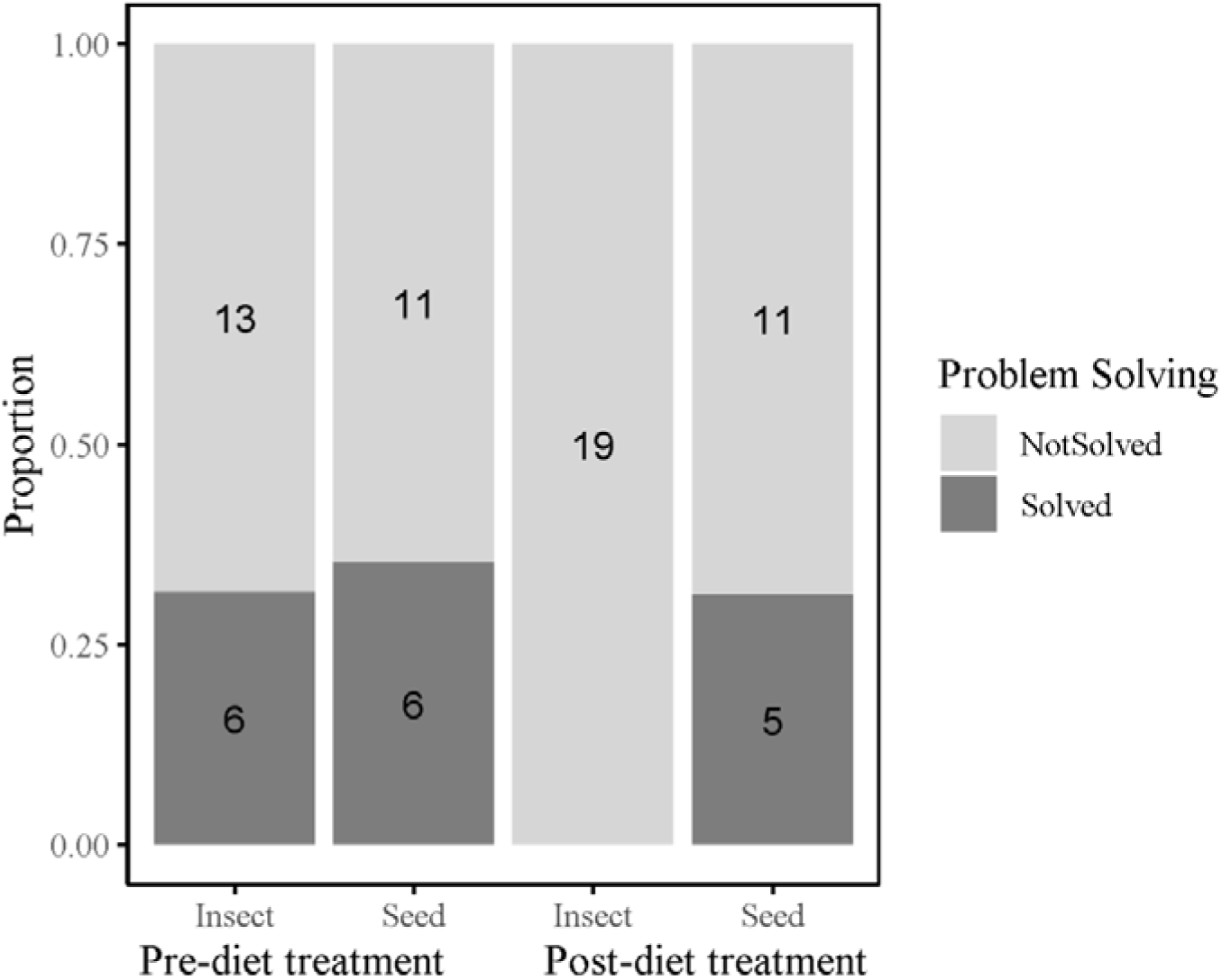
Problem solving performance (PSP) as a measure of innovation. Number of individuals that solved (dark grey) and number of birds that did not solve (light grey) pre- and post-dietary treatments.

## DISCUSSION

We demonstrate that an experimentally induced dietary change caused significant alterations to the gut microbiome diversity, and to phylum- and genus-level abundance in a wild bird species, which in turn was correlated with reduced innovative behaviour. To our knowledge, this is the first experimental evidence from a non-model laboratory organism that gross microbiota composition is associated with a behavioural trait, in this case PSP. Nevertheless, owing to the correlational nature of these relationships, it is possible the parallel changes to gut microbiome and behaviour occurred independently and were driven solely by the dietary treatment. We discuss the extent to which our results may point to a causal link between microbiota and behaviour, and their broader relevance to foraging ecology, the microbiome-gut-brain axis, and environment-behaviour interactions.

Seasonal and geographic differences in gut microbial communities in wild mammals have been attributed to changes in food availability ^21–23^. In a population of wild birds temporarily taken into captivity, we show that changes in microbial community composition are sensitive to dietary changes, independent of other factors that may differ with seasonality and impact on gut microbiota, such as hormonal differences ^50^, because the changes here were recorded under controlled conditions over a two week period. We show that phylum-level and genus-level diversity changes to the gut microbiome were caused by the diet, which may be attributed to differences in relative protein, fat and fibre content ^27,28^. At the phylum-level, we found higher abundance of Bacteroidetes present in subjects with lower fat, protein and fibre intake, a finding in line with Clarke et al ^28^; in contrast to that study, however, we did not find a decrease in Firmicutes, and found an increase in Proteobacteria. These differences in dietary effects on the gut microbiota may be due to differences in the microbiome between birds and mammals ^51^, and the type of fat and/or fibre ^reviewed in 52^. At the genus level, the insect diet caused a reduction in *Clostridium spp. –* taxa that have previously been reported to be positively associated with host health in Eastern bluebird chicks (*Sialia sialis*) supplemented with mealworms ^53^, and in poultry fed diets that differed in animal protein and in non-digestible non-starch polysaccharides ^25^.

Birds in the insect diet showed a decrease in Chao1 index, a measure of diversity which estimates taxa abundance and rare taxa missed from undersampling ^54^. This result may be because the insect diet itself was less diverse than the seed diet. While birds in both the seed and insect diets showed both decreases and increases in genus-level abundance, only birds given the insect diet showed significant changes in diversity and phylum-level abundance. The lack of change in the seed treatment could perhaps be explained because our birds had already been taking seeds at the feeders we used to lure them for capture, and because great tits consume a high proportion of plant-based foods in the winter ^55^. In urban and rural house sparrows, microbiota changes as a result of dietary manipulation were greatest when fed on diets that contrasted from their place of origin ^26^. In the current study, the use of garden feeders at all trapping sites could have precluded any effect the seed diet may have on the gut microbiome and/or PSP. Moreover, the use of garden feeders may have homogenised the diets of both urban and rural populations. Alpha diversity and urbanisation were positively associated in male white-crowned sparrows (*Zonotrichia leucophrys*) ^56^, but we found no such effect. There was, however, a significant difference in beta diversity as well as a higher proportion of Actinobacteria in urban birds compared to rural birds, similar metrics to those that have previously been shown to be related to urban environments in house sparrows (*Passer domesticus*) ^49^. Our experiment in captivity aimed to mimic variation in individual food consumption, or in seasonal food availability. Longitudinal sampling of individuals across seasons would be necessary to confirm whether similar microbial taxa changes found in this study would also occur under natural conditions. For example, the invertebrate species accessible in the wild would differ from those provided in our experiment. Whatever the reason for limited effects of habitat on the microbiome, our manipulation showed clearly that diet does modify the microbiome in this model species in field ecology, and that more generally environmental drivers of variation in diet may play a significant role in driving these effects.

Dietary manipulation in this experiment affected not just the gut microbiome but was also correlated to the likelihood of solving. Those on the insect diet had reduced microbiome diversity and were less likely to solve, suggesting a potential causal link whereby the dietary induced reduction in the microbiome may have impeded PSP via the microbiome-gut-brain axis. Our results showed that the indices of microbial community diversity that decreased as a consequence of diet (i.e. Chao1 index, beta diversity) were the same metrics that were associated with variation in PSP. Proteobacteria and Bacteroidetes increased following an insect diet, but these two phyla were not associated with PSP, nor were the genus-level microbial taxa that were altered as a consequence of diet, suggesting that the microbial community structure as a whole may be important for regulating behaviour. Moreover, our results suggest that less abundant taxa may contribute to this relationship because two of the metrics that were significantly associated with PSP were those that are most sensitive to differences in low-abundance features (i.e. Chao1 index and unweighted unifrac), although the significant relationship with Shannon’s diversity also suggests OTU evenness is an important predictor. We also note that while we detected these effects, our sample size was relatively low owing to logistical constraints associated with a short field season, the long duration birds were held in captivity, and limited holding capacity in captivity. While clinical studies on the microbiome-gut-brain axis typically use in the range of 6-12 animals per treatment, much less than we used here, microbiomes from wild animals are likely far more variable ^13,57^, and therefore further studies with even larger sample sizes will likely be necessary to confirm the results we present here.

Motivation can influence PSP ^reviewed in 58^ and an insect diet may have decreased motivation to engage in the problem solving task baited with an insect reward. However, the insect diet did not influence the birds’ motivation to consume the freely available wax worm, irrespective of dietary treatment; and birds solved on day one when other insects (i.e. mealworms) were freely available, indicating that wax worms are a highly-valued and preferred food reward. Furthermore the links between PSP and the microbiota were significant, even after controlling for dietary manipulation as a fixed effect (see Table S2), and for whether birds ate the freely available waxworm. Nevertheless, we cannot discount the possibility that birds were motivated to meet specific caloric or nutrient requirements due to differences in diets, although there is evidence that the gut microbiome can mediate host diet selection ^59^. Diet may have also affected how motivated the birds were to work for the waxworm reward inside the apparatus. PSP was measured as a binary response, and birds may have differed, for example, in whether they tackled the task at all, and how persistent they were at attempting to solve the task. PSP is a composite trait involving motivation, exploration, persistence and associative learning ^58^, many of which have been previously reported to be associated with the microbiome-gut-brain axis ^7,60,61^. It is not clear which of these traits was influenced, whether it was mediated directly by the diet, by the change in the gut microbiome, or a combination of both. Identifying causality in microbiome-mediate behaviour in wild animals remains a challenge particularly because environmental inputs such as diet and stress can either act independently on behaviour, or together with the gut microbiome ^13,37^. Here we ruled out stress as a determining factor as baseline and stress-induced corticosterone did not explain PSP. Further manipulative experiments, such as using a wide range of food rewards, or microbial transplantation experiments ^13,62^ in the absence of dietary differences would provide additional support that the microbiome itself caused the change in problem solving performance.

How the gut microbiome impacts behaviour via the gut-brain axis may be attributed to the metabolic functions of the microbial community itself ^e.g. 12^, or derived from the diets of the host ^reviewed in 37,63^. Studies have attempted to disentangle nutritional and microbial effects on behaviour by depleting the original microbes of the hosts and re-introducing specific bacterial organisms, or transplanting gut microbiomes between hosts ^e.g. 61,64^. And while we did not manipulate microbiome directly owing to logistical and ethical constraints, the aim of our study was to test diet-microbiome-behaviour relationships within an ecologically relevant context that would translate to wild animals in their natural environment. To control for nutritional deficiencies that may have an impact on behaviour independent of microbiome, we provided vitamin supplements, though other nutritional differences cannot be discounted. Fat content and fibre content was five-fold and three-fold higher in the seed diet compared to the insect diet, respectively. Mice fed on high fat diets, or given microbiome transplantations from obese donors show poorer cognitive performance than control mice ^8,61^; whereas, our study showed that birds fed the lower-fat diet (i.e. insect) had poorer PSP. Having a higher proportion of fibre present in the seed diet may have offset any negative effects of a high-fat diet. Non-digestible carbohydrates are fermented by gut microbes in the large intestines, promote the growth of microbial organisms and can have positive effects on cognition and behaviour in mammals ^reviewed in 65^. Metabolomics profiling would be an informative future endeavour to provide a functional assessment of microbial products such as short chain fatty acids involved in gut-brain axis communication ^reviewed in 12,65,66^ to pinpoint further the mechanisms underlying phenotypic plasticity in innovative behaviour.

Contrary to our predictions, we did not find that individual variation in PSP was positively associated with natural variation in microbial alpha diversity, as measured on the first day in captivity and before dietary manipulation. Instead, beta diversity was dissimilar between solvers and non-solvers, although this result was marginally non-significant and correlational. One reason for the lack of an effect is that all of the birds were feeding on the same food at the feeders, and individual variation in foraging on other food sources may simply not have been pronounced until we manipulated the diet in captivity. Further manipulative investigations are needed to pinpoint causal directions of these relationships under natural conditions, in particular whether innovative behaviour leads to variation in microbial diversity through food access, or indeed whether innovation arises because of microbial diversity caused by some other mechanism, such as developmental effects related to habitat quality, as previously shown in this species ^45^. Together our findings lend support to the hypothesis that the gut microbiome, innovation and diet are interlinked, and more generally that the microbiome may be an important source of variation in ecologically important behaviours, especially those linked to diet and foraging.

## MATERIALS AND METHODS

### Subjects

Thirty six great tits were captured and held in captivity between January 10^th^ and March 9^th^ 2017 across four sites. Two sites were within Cork city (urban), 1.6 km apart (51°54’32.4”N 8°28’12.0”W and 51°53’56.1”N 8°29’15.4”W), and two were in deciduous woodlands (rural) 23 km apart (51°43’08.4”N 8°35’52.8”W and 51°51’37.5”N 8°47’01.1”W), and located at least 23km from the urban sites. We alternated between trapping sites throughout the experiment, with the following exceptions owing to temporal and spatial overlap of independent experiments: we stopped trapping at one rural site on January 22^nd^, and commenced trapping at one urban site on the 22^nd^ February. Birds were lured using feeders containing sunflower seeds and peanuts. All birds were banded with rings issued by the British Trust of Ornithology for individual identification. Birds were sexed (male/female) and aged (adult/juvenile) from plumage characteristics (i.e. breast stripe size and presence of moult limits) ^67^. Juveniles refer to birds hatched the previous calendar year (i.e. spring 2016). Upon capture, birds were transported to the aviary facilities at University College Cork and singly-housed in wire cages (45 × 50 × 60 cm) containing two wooden perches and were visually isolated from each other.

### Faecal sampling

Fresh, undisturbed faecal samples were collected within 1 hour of arrival into the aviary, and again on Day 13 of captivity. A clean sheet of brown paper was placed on the floor of each cage for faecal collection. DNA concentration extracted from bird faecal samples is substantially less than from mammalian faeces (personal observation), therefore to improve DNA yield, we used the paper to soak liquid urea away from the faecal matter as urea can act as a downstream inhibitor to amplification ^68^. Using sterile inoculation loops, we transferred the faecal matter into tubes containing 1ml of 100% ethanol and stored tubes at -20 degrees Celsius. We recognise that the paper, as well as the wire cages open to the air may have been sources of contamination, but this is not expected to systematically bias our results as our experimental design was fully counterbalanced.

### Dietary manipulation

From day 2-13 of captivity, birds were given one of two different dietary treatments: Seed, n = 17; and insect diet, n = 19. These diets were designed to reflect ecological variation seen in the wild, for example changes in the availability of seed or animal food sources ^55,69^, or perhaps reflecting potential individual differences in dietary specialisations ^70^: The insect diet consisted of wax moth larvae (*Achroia grisella*) and mealworm larvae (*Tenebrio molitor*). Notably, these two diets differed in the relative content of protein, fat and fibres (Table S1), features of diets that have previously been associated with differences in host microbiota in mammals ^52^. Mealworms were provided ad libitum, and five wax worms were provided each morning and each evening (except during the problem solving task). The seed diet consisted of sunflower hearts, peanuts and suet. We provided birds in the seed diet five mealworms and one wax worm on day seven of captivity for welfare reasons as we did not want to deprive the birds of a preferred food item as preference for mealworms/wax worms was observed in our population (pers. obs), and on the basis of other work ^e.g. 71^. To limit more general nutritional deficiencies, all birds received vitamin powder mixed with their food and drops mixed in their water (AviMix). Birds were assigned to treatment groups randomly, counterbalanced for age, sex and habitat.

### Problem solving assay

To quantify individual innovative problem solving performance birds were presented with a foraging task, derived from a similar foraging task ^40,41,43^ in which birds are required to pull a stick to release a platform holding a food reward. The current task also included two alternative solutions designed to control for potential differences in motor skills among individuals – a door that could be pushed to the side, and a string attached to a food reward, however, none of the birds solved the latter method. Following ^40,41,43^, the birds were given the task overnight from one hour before sunset to two hours after sunrise, times when birds were most active and motivated to forage. This same apparatus was presented once on Day 1 of captivity and once on Day 12 of captivity. Due to the length of the trial, birds were not food deprived for welfare reasons. During the first trial, all birds had access to mealworms, peanuts and sunflower hearts ad libitum. During the second trial, birds had access to their assigned diets ad libitum. During both trials, wax worms, a highly preferred food reward ^43,72; G. Davidson personal observation^, were placed inside a transparent Perspex tube 16cm (height) x 5cm (width). The worms could be accessed by solving at least one of three solutions: 1) by pulling a lever to drop a platform holding a worm; 2) by pushing a door to the side; and 3) by pulling a string attached to one of the worms from the top of the tube. By having multiple access possibilities, PSP could be assessed without limiting solutions to one particular motor action that may be more feasible in some individuals over others. While we report the numbers of each solving method in the PSP results, there was not sufficient power to analyse these specific differences statistically. Moreover, our aviary setup did not facilitate video recording, therefore PSP was recorded as a binary variable (i.e. solved versus not solved). At the start of the problem solving assay, a freely available wax worm was placed outside, at the base of the problem solving device to measure birds’ motivation to approach the apparatus and consume the wax worm. Wax worms were otherwise not provided when the problem solving task was presented. One bird died of unknown causes following 10 days of captivity and therefore only data from Day 1 were included for this bird.

### Glucocorticoid assay

We collected blood samples at capture (day 1), and immediately after the second problem solving task (day 13) to quantify baseline and stress-induced corticosterone as part of an additional study (unpublished, in preparation). We used this data to test whether problem solving performance could be explained by stress levels.

### Microbiome analysis

Microbial DNA was extracted using the Qiagen QIAamp DNA Stool Kit, following the “Isolation of DNA from Stool for Pathogen Detection” protocol with modifications described in ^73^ to increase DNA yield and remove excess inhibitors expected to be present in the uric acid of bird faeces [but see ^74^ where they found no evidence of uric acid in faecal matter from a subset of avian species]. A 0.10 - 0.20 g aliquot of each faecal sample was added to the kit, alongside a negative control.

The V3-V4 variable region of the 16S rRNA gene was amplified from the DNA extracts using the 16S metagenomic sequencing library protocol (Illumina) as described in Fouhy *et al* ^75^. In the current study, each PCR reaction contained 23 µl DNA template, 1 µl forward primer (10 µM), 1 µl reverse primer (10 µM) and 25 µl 2X Kapa HiFi Hotstart ready mix (Roche, Ireland), to a final volume of 50 µl. Two negative controls were run in parallel – one from the DNA extraction and one containing PCR water instead of DNA template. Of the bird samples, ten failed to amplify and were not pooled for sequencing (Day 1: n = 1 (seed), Day 13: n = 3 (seed), 6 (insect)). Successful PCR products were cleaned using AMPure XP magnetic bead based purification (Labplan, Dublin, Ireland). Samples were sequenced at the Teagasc Sequencing Centre on the MiSeq sequencing platform, using a 2 x 300 cycle kit, following standard Illumina sequencing protocols.

Three hundred base pair paired-end reads were assembled using FLASH (FLASH: fast length adjustment of short reads to improve genome assemblies). Further processing of paired-end reads including quality filtering based on a quality score of > 25 and removal of mismatched barcodes and sequences below length thresholds was completed using QIIME (Quantitative Insights Into Microbial Ecology) version 1.9.0 ^54^. Denoising, chimera detection and clustering into operational taxonomic units (OTUs) (97% identity, following Konstantinidis & Tiedje ^76^) were performed using USEARCH version 7 (64-bit). OTUs were aligned using PyNAST (PyNAST: python nearest alignment space termination; a flexible tool for aligning sequences to a template alignment) and taxonomy was assigned using BLAST against the SILVA SSURef database release version 123. Alpha diversities (observed OTUs, Chao1 and Shannon’s index) were generated from all OTUs in QIIME.

### Statistical analyses

The QIIME files (OTU table, taxonomic table, phylogenetic tree file) and metadata files were analysed using phyloseq ^77^ in R Statistical Software ^78^. Singletons and taxa present at <0.005% were removed ^79^. We applied cumulative sum scaling normalisation to standardise library size across samples for beta diversity analysis. We included all taxa in the dataset, most notably those that are often viewed as ‘dietary contaminants’ (i.e. microbes originating from ingested food, such as cyanobacteria present in plants). This is because not all cyanobacteria are dietary contaminants ^80^ and because our study specifically tested differences between diets. Thereby removal of one so-called ‘dietary contaminant’ specific to plants (i.e. cyanobacteria), but none specific to insects (e.g. insect microbiome) could systematically bias our results. We also note that the non-sterilised water was an additional source of dietary contaminants, although this would not systematically differ between dietary treatment groups.

Linear Mixed Models (LMMs) and Generalized Linear Mixed Models (GLMMs) were run using lme4 ^81^, and where relevant, p-values were obtained using lmerTest ^82^ in R version 3.5.2 ^78^. Terms with p > 0.1 were sequentially removed from the models, starting with interaction terms. Site ID and Bird ID were included as random effects. Response variables were transformed where necessary to meet assumptions of normality (Natural log (Ln) or square root transformed), and therefore all models were run with a Gaussian distribution unless stated otherwise.

### Dietary effects on microbiome

The effect of diet on the relative abundance (i.e. percentage abundance relative to all other phyla) was tested for Firmicutes, Proteobacteria, Bacteroidetes, Tenericutes and Actinobacteria as these were the most abundant phyla. Proteobacteria was modelled as a proportion and run with a binomial distribution, as it could not be transformed to fit a normal distribution. Three different measures of alpha diversity were used as response variables to test the effect of diet on the microbiome community: Shannon’s index, total observed OTUs, and Chao1. The above gut microbiome metrics did not differ between pre-assigned dietary groups (supplementary materials, Figure S1), therefore we used diet as a three level factor as a fixed effect: Diet (pre-dietary assignment (day 1), insect (day 13) and seed (day 13)). Habitat (rural, urban), age (juvenile, adult ^67^) and sex were included as fixed effects. To avoid overparameterization of models, we tested the interaction between diet and habitat only, as we expected inherent differences in diets between birds from the two habitats. Sampling date (i.e. days since January 1^st^ did not predict alpha diversity or phylum-level abundance therefore we did not include this term in all models. We also investigated the effect of diet on genus-level relative abundance with mixed regression models using the web-based software Calypso (version 8.72) ^83^. ID was included as a random term and significant taxa were adjusted by a false discovery rate (FDR) whereby p-values of less than 0.05 were considered statistically significant.

### Dietary effects on problem solving performance

PSP was modelled as a binomial distribution (solved vs not solved) with diet (seed, insect), experiment day (day 1, day 12), habitat (rural, urban), age (juvenile, adult) and sex included as fixed effects. We included interactions between diet and habitat type and diet and experiment day. To test whether diet affected motivation, we ran a binomial GLMM with consumption of the freely available wax worm (Yes/No) as the response variable, assuming that birds who took the waxworm were more motivated to solve than those who were not. The fixed and random effects were included as described for the problem solving analysis above.

### Relationship between problem solving and microbiome

We tested whether natural variation in the gut microbiome correlated with PSP on Day 1. We tested for associations between microbial community and PSP as a binomial response variable (solved vs not solved) in GLMMs for each microbial community measurement (the top five phyla, and the three measures of alpha diversity), and included habitat, sex and age as fixed effects and site as a random effect. Beta diversity was calculated in three ways: Jaccard distance matrix, and phylogenetic weighted (accounting for relative abundance of taxa) and unweighted (presence/absence of taxa) unifrac distance matrices. Each matrix was analysed using permutational multivariate analysis of variance (ADONIS) with 1000 permutations. We tested for homogeneity of dispersion, and found there was no difference in dispersion between groups. We also investigated the relationship between problem solving and relative abundance at the genus-level as described above. To test whether gut microbiome alteration as a consequence of dietary manipulation caused a change in PSP, we ran the same analyses described above, but included data from Day 12 and diet as a fixed effect with bird ID as a random term.

### Research and Animal Ethics

This study was conducted under licences from the Health Products Regulatory Authority (AE19130_P017), The National Parks and Wildlife Services (C11/2017) and the British Trust for Ornithology, and permission from Coillte Forestry and private landowners. The research project received ethical approval from the Animal Welfare Body at University College Cork, and was in accordance with the ASAB (Association for the Study of Animal Behaviour) Guidelines for the Treatment of Animals in Behavioural Research and Teaching.

## Acknowledgments

James Nichols, Ally Phillimore and Sarah Knowles for helpful discussions on molecular methodology; Shane Somers for advice on statistics; Jennifer Coomes for assistance with fieldwork; Dr. Paul Cotter, Dr. Fiona Crispie and Ms. Laura Finnegan from the Teagasc Sequencing facility for their role in relation to the 16S rRNA sequencing. We thank three anonymous reviewers for their valuable comments. Funding for GLD, ACC, MSR, IGK, IDH and JMSC from the European Research Council under the European Union’s Horizon 2020 Programme (FP7/2007-2013)/ERC Consolidator Grant “Evoecocog” Project No. 617509, awarded to JLQ. This work was funded in part by Science Foundation Ireland through APC Microbiome Ireland.

## Data accessibility

Our data will be deposited at Dryad, and sequence data uploaded to a repository such as NCBI Sequence Read Archive, if the manuscript is published.

## Authors contributions

GLD, ACC and JLQ designed the experiment with input from CS and RPR. GLD and ACC ran the experiment with assistance from MSR, IGK, IDH and JMSC. GLD, NW and CNJ carried out DNA extraction and library prep. FF and GLD analysed data. GLD wrote the manuscript with input from all authors. All authors gave final approval for publication.

## Additional Information

The authors have no competing interests.

## Legends

Table 1. Means and se of the most abundance phyla across treatment groups

Table 2. Summary of the relationships between microbiota and host traits. Significant relationships are shown as either positive (+), negative (-) or as p-values (i.e. beta-diversity where differences do not represent directionality). NS = Non-significant.

Figure 1. Relative abundance of top seven phyla across dietary treatments. Proteobacteria and Bacteroidetes significantly increased following the insect diet, but not the seed diet.

Figure 2. Relative abundance of bacterial phyla per sample for birds before and after their dietary assignments of either insect diet (ID) or seed diet (SD).

Figure 3. Alpha diversity for birds pre-dietary assignment (Day 1), and following the insect diet (ID) and the seed diet (SD) for a) Chao1 (sqrt), b) Shannon’s index, c) Observed OTUs (Ln). Coloured points denote individual data points, black points and line denote mean and ± SE. p-values represent the comparison between post-insect diet group and day 1.

Figure 4. Nonmetric Multidimensional Scaling (NMDS) ordination plots derived from beta diversity measurements for (a) weighted unifrac distances of diet (Day 1, insect diet (ID) and seed diet (SD), and (b) unweighted unifrac distances of problem solving performance (not solved, solved). Ellipses represent standard deviations around the centroids of the groups. Numbers in brackets refer to the variance explained by NDMS axes.

Figure 5. Natural variation in gut microbiome diversity and problem solving performance for a) beta diversity (Nonmetric Multidimensional Scaling (NMDS) ordination plots), and b) Shannon’s index. Coloured points denote individual data points, black points and line denote mean and ± SE.

Figure 6. PSP-alpha diversity relationships in a) Chao1 (sqrt), b) Shannon’s index, c) Observed OTUs (Ln) including data points from both day 1 and day 12. Coloured points denote individual data points, black points and line denote mean and ± SE.

Figure 7. Problem solving performance (PSP) as a measure of innovation. Number of individuals that solved (dark grey) and number of birds that did not solve (light grey) pre- and post-dietary treatments.

